# Mortality and coexistence time both cause changes in predator-prey co-evolutionary dynamics

**DOI:** 10.1101/2020.04.08.031146

**Authors:** Thomas Scheuerl, Veijo Kaitala

## Abstract

All organisms are sensitive to the abiotic environment, and in multispecies communities a deteriorating environment increasing mortality and limiting coexistence time can cause ecological changes. When interaction within the community is changed this can impact co-evolutionary processes. Here we use a mathematical model to predict ecological and evolutionary changes in a simple predator-prey community under different mortality rates and times of coexistence, both controlled by various transfer volume and transfer interval. In the simulated bacteria-ciliate system, we find species densities to be surprisingly robust under changed mortality rates and times both species coexist, resulting in stable densities. Confirming a theoretical prediction however, the evolution of anti-predator defence in the bacteria and evolution of predation efficiency in ciliates relax under high mortalities and limited times both partners interact. In contrast, evolutionary trajectories intensify when global mortalities are low, and the predator-prey community has more time for close interaction. These results provide testable hypotheses for future studies of predator-prey systems and we hope this work will help to bridge the gap in our knowledge how ecological and evolutionary process together shape composition of microbial communities.

## Introduction

The composition of microbial communities is sensitive to the abiotic environment [1], which changes growth of individual species [2,3] and the interaction with other community members [4,5]. Abiotic modifications can affect communities in several ways, such as destabilization of predator-prey systems [6] and a stable predator-prey community may collapse when a keystone is lost [7]. This may happen, when there is not enough prey to maintain the predator resulting in extinction events [8]. Thus, a change in the abiotic environment may disrupt predator-prey systems and destabilize entire communities [6,9]. Following this, environmental changes can affect community structure and composition and disrupt vital functions pivotal for ecosystem functioning. Such triggering changes in the abiotic environment may include the use of antibiotics [10], or eutrophication of lake ecosystems [11].

A common effect of environmental change is the modification of the mortality rate [12] and for how long the community can grow without further disturbance. These two aspects can be easily implemented in laboratory experiments. In fact, a standard method using microbial communities to study predator-prey dynamics involves periodic transfer of the cultivation to fresh media [13–16], which removes a large proportions of the population at specific time points to restart the growth cycle with fresh nutrient conditions. The volume transferred, and the time point of transfer therefore determines mortality and coexistence time between interaction partners.

In liquid media containing all nutrients for rapid cell division, microbes can grow extremely quickly, which makes them suitable study organisms for experiments exploring ecological and evolutionary dynamics over several generations [17]. This, however, means that populations reach limiting conditions quickly. To keep the growth conditions constant, populations are either maintained in chemostat systems [18–20], or transferred to fresh conditions regularly (commonly every 24 and 72 hours) [4,13,16,21,22]. Transferring a small part of the populations every few days is a classical approach to keep populations constantly growing and to avoid growth plateaus, e.g. reaching carrying capacity, once nutrient limitation occurs [23]. This is an approach we focus on in this work. The two key parameters, *transfer volume* and *transfer interval*, are often chosen without further investigation, but the underlying mortality rates and coexistence times may have profound effects on the observed results.

Theoretically, increasing global mortality may result in situations where the evolution of defence and attack traits should decrease because of global weaker interaction strength and the predator may be lost due it’s general lower abundance compared to prey organisms [8,24]. Contrary, extending the time both partners coexist, at least theoretically, should stabilize predator-prey coexistence up to a point where predators overexploit the prey and intensify evolutionary traits determining interspecific interaction. Missing in our knowledge is how both factors, global mortality and time of coexistence, together affect ecology and evolution in a predator-prey community. Experimental tests of ecological and evolutionary dynamics in microbial predator-prey systems are extremely laborious and applying more than one transfer volume and transfer interval quickly gets non-feasible. Theoretical modelling offers a convenient approach out of this dilemma.

Here, we present such an approach taking advantage of a mathematical model simulating microbial predator-prey dynamics [25]. A recent experimental study tracked interaction between *Pseudomonas fluorescens* bacteria as prey and *Tetrahymena thermophila* ciliates as predators, to investigate ecological and evolutionary dynamics [14]. In the experiment, bacteria and ciliates were maintained in nutrient rich King’s B medium and 1% of the community was transferred to fresh conditions every 48 hours. Using the data of this study we previously built a mathematical model [25] and simulated the observed predator-prey dynamics, but kept the parameters *transfer volume* and *transfer interval* the same as in the experimental study [14]. This model allows us to track ecological dynamics in form of predator and prey densities and reveals how prey species evolve anti-predator defence traits and how predator species evolutionary increase their predation efficiency.

In this paper we simulate the effect of alternative transfer volumes (mortality) and time intervals to the next transfer (time of coexistence) to study how mortality and time of coexistence affect ecological and evolutionary dynamics. Consider population growth curves of bacteria, as prey and ciliates as predators for a single growth period (Fig. 1). Bacteria will start growing exponentially until they reach carrying capacity (Fig. 1a). When bacterial densities are high enough the ciliates will consume the bacterial cells and will increase in density (Fig. 1b); this way reducing bacterial densities until ciliates can grow no more due to lack of prey. It can be easily seen that time of coexistence (transfer interval) and mortality rate (transfer volume) can have major impact on the next growth period. If the coexistence time (transfer interval) is short, only bacterial densities may be high and ciliate densities may still be neglectable. If the transfer interval is long, ciliates may have already consumed most bacteria and the next growth cycle is initiated with ratios resulting in extreme changes. The transfer interval determines the ratio between prey and predator for the next growth period. If global mortality is high (small transfer volume) the time both partners interact is shorter because initial densities are lower and encounter rate is reduced, which may mean reduced strength of natural selection, whereas a low mortality may place both partners in close contact from the beginning of the experiment and intensify interaction.

**Figure 1.**
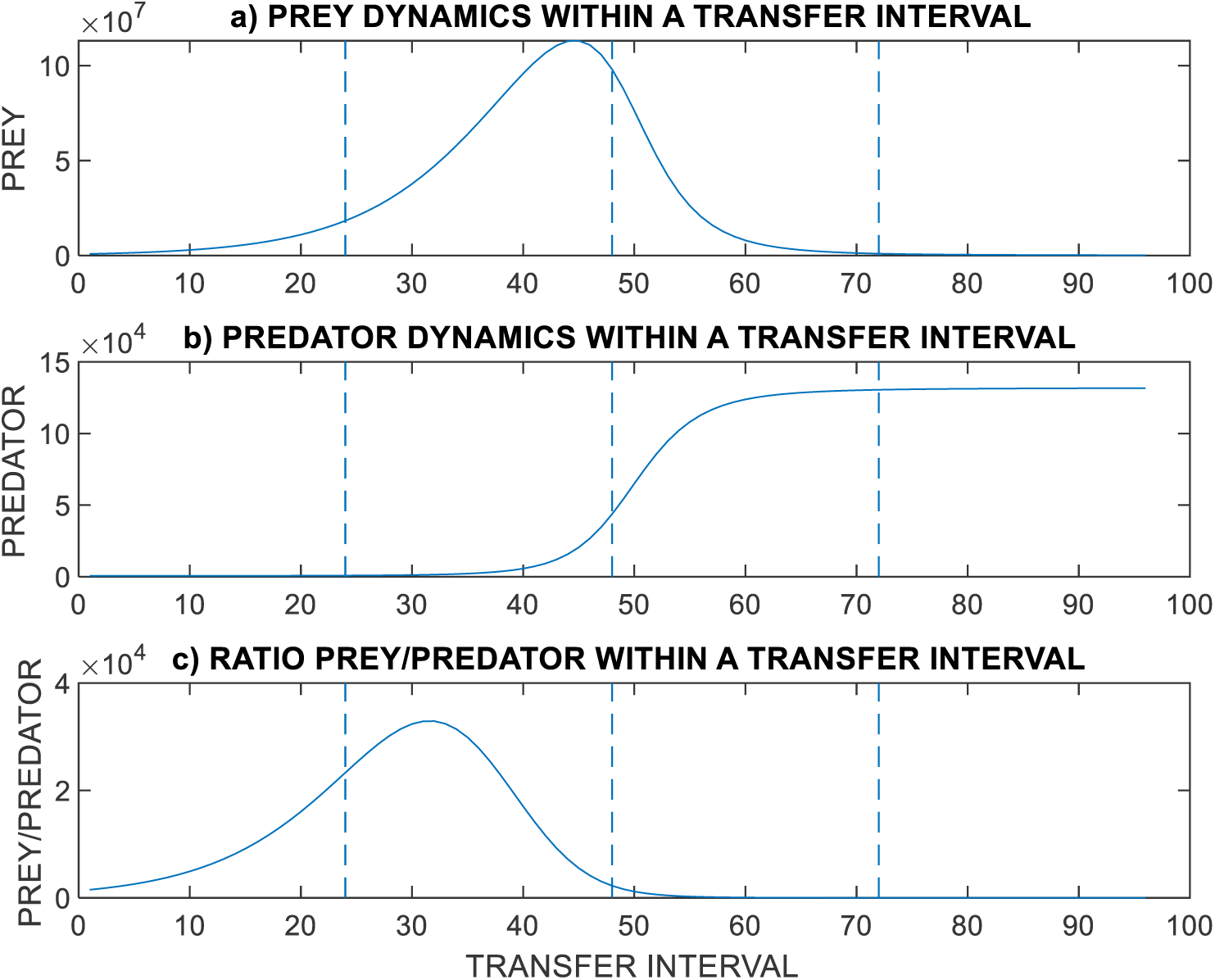
Hypothetical example dynamics of a predator-prey system within a sampling period. a) Prey abundance. b) Predator abundance. c) The ratio of the prey and predator abundances. Three alternative transfer intervals are indicated by vertical lines: 24, 48, and 72 hours.

Expanding our previous model [25] we report how global mortality and time of coexistence affect predator-prey communities and expand the prior literature by exploring scenarios impractical in experimental studies. Our theoretical findings suggest that both, mortality and coexistence time, have effects on the community. First, increasing the mortality rate we find that coexistence is threatened. The predator may be lost, and evolutionary rates decrease. In contrast, decreasing mortality maintains coexistence but destabilizes initial ecological dynamics, while the adaptation of predator and prey increases. Second, decreasing time of coexistence has similar effects driving populations extinct and decreases evolutionary rates. Increasing the time of coexistence destabilizes initial ecological dynamics and intensifies evolutionary rates. Interestingly, if no extinction happens, the community dynamics are predicted to be overall stable, indicating that predator-prey dynamics are rescued by evolution. According to this, we suggest that slight modification in mortality rate and transfer interval will have initial ecological effects, but extended effects on evolutionary measures.

## Results

### Stability of predator-prey communities and evolutionary dynamics

We estimated parameters necessary for our model using data presented in a study exploring ecological and evolutionary dynamics in a life bacteria-ciliate system [14]. This experiment maintained the organisms using a 99% mortality rate (1% transfer) and a coexistence time of 48 hours (3.3 generations) before starting the next growth cycle. The experimental data and our model predictions consistently result in coexisting prey and predator populations under these conditions. Prey densities increase over time because anti-predatory traits evolve and bacteria get less eatable by ciliates [14]. The predator densities decrease over time as prey gets better defended against attack. Coevolution of the predatory attack rates prevents further decrease in the predator densities [25] and the community stabilizes after a few transfers. This suggests arms-race dynamics [26] where the defence and attack traits evolutionarily increase during this experiment [25].

### Changing global mortality affects initial dynamics and evolutionary rates

To explore how increased mortality affects predator-prey communities we successively modified the transfer volume in our model (Fig. S1) but kept the transfer interval constant at 48 hours. An increased mortality may result in a change in ecological dynamics (e.g. extinctions) and in reduced rates bacteria and prey encounter each other (at least for the initial growth period), which may relax anti-predator and attack rates in bacteria and ciliates respectively.

Transferring only 0.3% of the populations (compared to 1% as in the original study), thus increasing the mortality up to 99.7%, results in extinction of the predator. This is the predicted transfer volume where only the prey can survive in this system. Probably, the mortality rate is so high at this point that there is too little prey available and predators are unable to catch enough food to grow rapidly enough to compensate mortality. While there is most likely enough prey available (we see 1.5 × 10^6^ bacterial cells per ml) these conditions obviously out-dilute the ciliates and they cannot compensate the loss via growth. Bacteria are dwindling towards extinction at a mortality rate of 99.9% (0.1% transfer volume). Considering these high mortality rates, the system, however, seems rather stable for experimental modifications. Decreasing the mortality below 99% (e.g. transfer volumes >1%) has surprisingly little effect (Fig. 2). When mortality is less severe, and the next growth cycle is started with higher densities, the initial dynamics seem to destabilize a bit. However, after a few transfers the fluctuation in predator-prey densities fade away and there is no obvious difference between 99% and 98% (or lower) mortality rates (Fig. S2).

**Figure 2.**
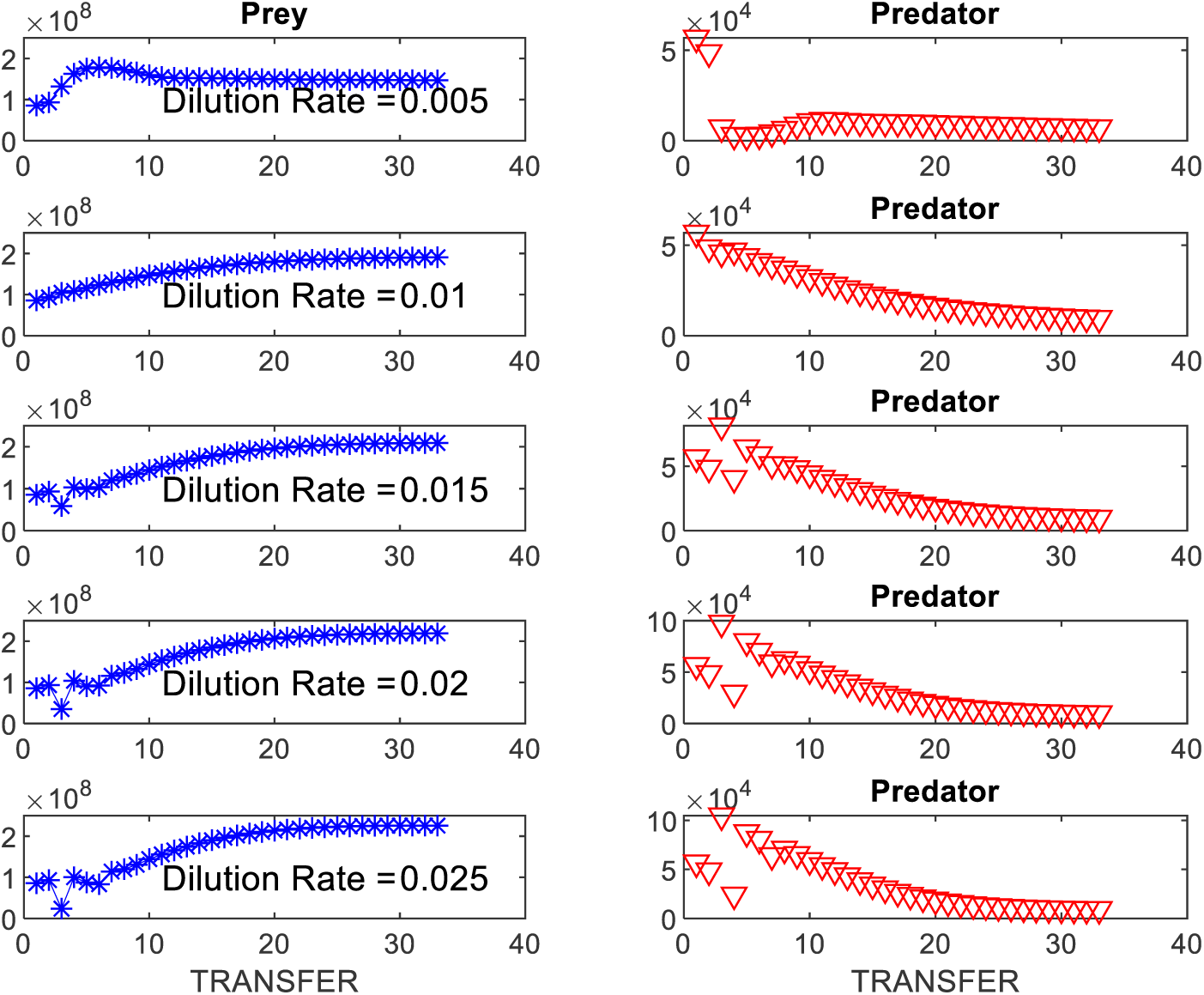
The effect of modifying the mortality rate on population densities controlled by applying different transfer volumes. The time of coexistence is kept constant at 48 hours. There are 33 transfers. The mortality rate is ranging between 99.9% and 97.5%. Bacteria (prey) is denoted by blue stars, the ciliates are represented by red triangles.

High mortality rates should release prey form predation pressure. This should result in a decrease of anti-predator defence evolution in bacteria, but an increase in predation ability in the ciliates. Indeed, our results indicate a change in the evolutionary rates. Our model successively predicts that bacteria evolve less anti-predator defence with increasing mortality (Fig. 3a). At high mortalities (low transfer volume) the anti-predator prey trait *u* only changes moderately, but when mortality is low (high transfer volume) we see a great change in evolution. On the ciliate side, we see a higher change in predator trait *v* (Fig. 3b and c) and the attack rate *a* under high mortalities as we would expect when predators are selected for higher efficiency due to reduced encounter events. The conversion rate *b* decreases over course of the experiment, but less under lower mortalities (Fig. 3c). At the extreme high end of mortality, when only the bacteria survive, anti-predator defence traits stop evolving (Fig. S2).

**Fig. 3.**
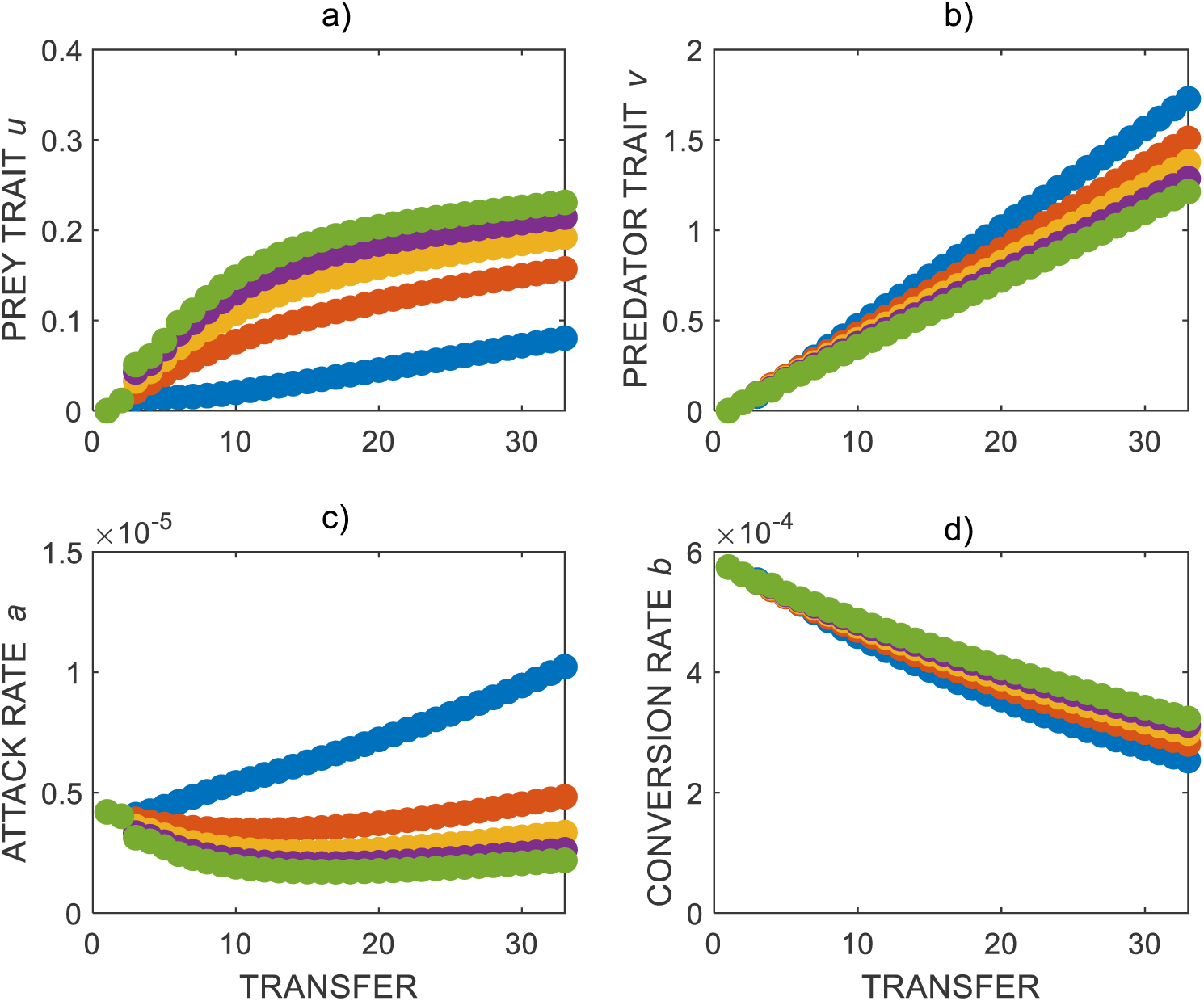
Evolutionary trajectories for bacteria and ciliates under different mortality rates. a) prey trait *u* defining the anti-predator defence level, for the ciliates the following parameters are b) predator trait *v*, c) predator attack rate *a* and d) predator conversion rate *b*. Dots in blue, red, yellow, magenta and green denote increasing mortality rates (dilution rate) with 0.5%, 1%, 1.5%, 2% and 2.5%, respectively.

Interestingly, after around 25 transfers, our model predictions are that the prey-predator ratios are the same for all mortality rates (Fig. S3). Before this happens, we see great differences in the bacteria-ciliate ratios with much more bacteria at highest mortality rate. At the highest mortality rate, predators need a prolonged time to catch up and to establish stabile populations. The final ratio, however, seems to be robust against different mortality rates.

### Changing time of coexistence affects ecological and evolutionary dynamics

Because we saw an effect of mortality on ecological and evolutionary dynamics in this system we next asked ourselves if the period both, prey and predator, grow together (transfer interval) may have an effect as well. As indicated in Fig. 1, unlike global mortality which keeps ratios sustained, this should affect the bacterial-ciliate ratio transferred to the next growth cycle. On the ecological side, this means that the transfer interval modifies the ratio between bacteria and ciliates for the next growth cycle which may affect timing when ciliates start to consume bacteria. On the evolutionary side, anti-predator and attack rates are expected to intensify under longer antagonistic interaction periods.

Applying different times of coexistence (transfer intervals) indeed resulted in various ecological dynamics (Fig. 4). In the original case, the populations grow for 48 hours before they are transferred to fresh conditions, which results in stable bacteria densities. The ciliate densities first steadily decrease but stabilize towards the end of the experiment at low densities. When the transfer interval is reduced to 36 hours the ciliates rapidly go extinct. At even lower transfer intervals (24 hours) the bacteria also cannot exist any longer and get out-diluted. Increasing the growth period has little effect on the ecological dynamics. Interestingly, the initial dynamics destabilize at longer times of coexistence. This may result from a change in the ratio after 48 hours, where predators already start to reduce prey densities significantly. This may mean, when the next growth cycle is initiated, proportional more predators are present. This effect however disappears at later growth cycles.

**Fig. 4.**
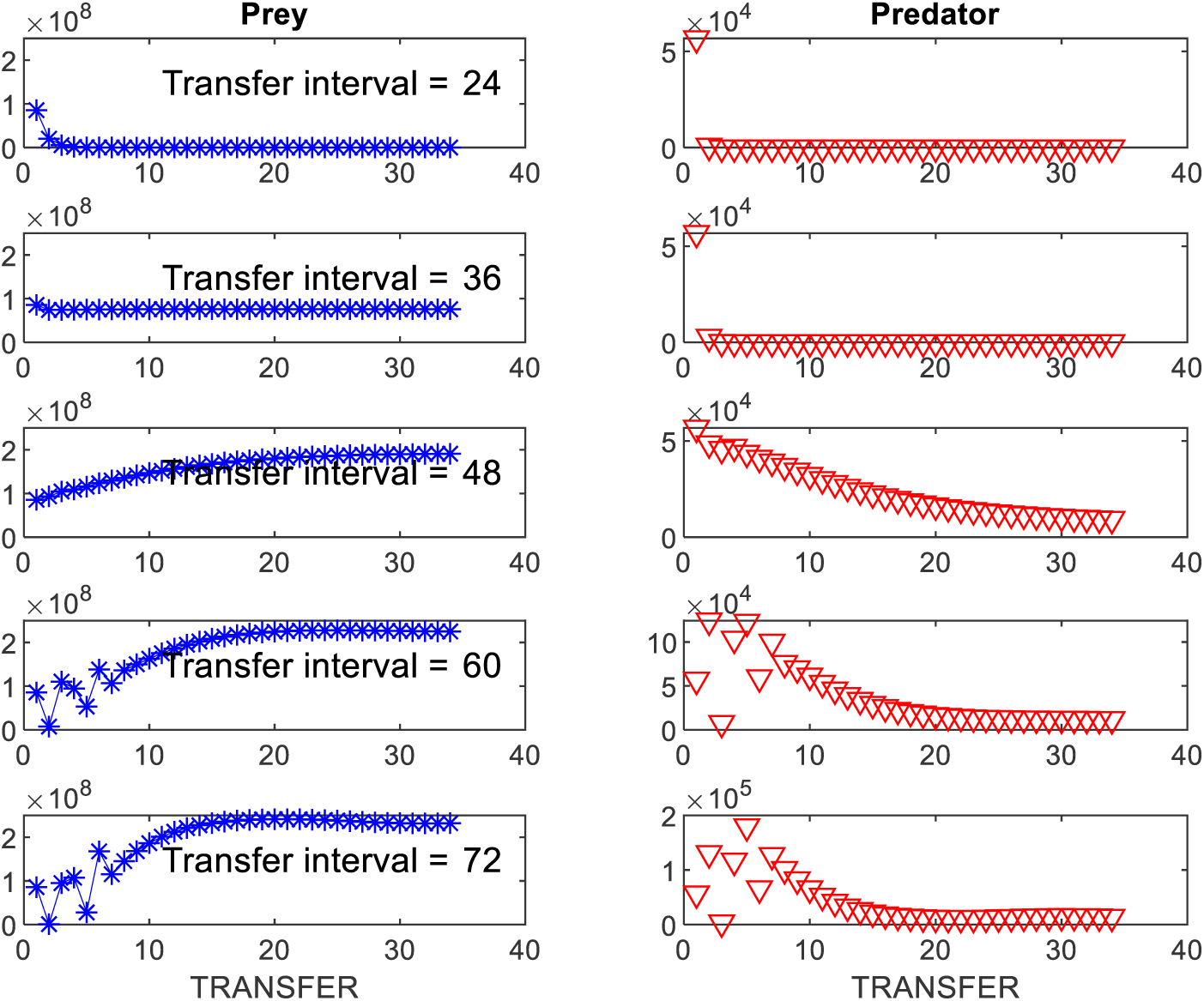
Effect of time of coexistence (transfer interval) on predator-prey dynamics. Bacterial densities (blue) and ciliate densities (red) are presented for different transfer intervals ranging from 24-72 hours. The transfer volume was constant with 1% every transfer. There are 33 transfers.

With increasing time bacteria and ciliates coexist during growth cycles we would expect the interaction to intensify, whereas when the coexistence time is limited any interaction may be weakened because low initial densities and reduced encounter rates. A transfer interval less than 48 hour quickly drives ciliates populations into extinction (Fig. 4) and evolution of anti-predator and attack rates in bacteria and ciliates hold (Fig. 5). Longer coexistence times than 48 hours result in increasing evolution of anti-predator trait defence trait *u* in the bacteria (Fig. 5a). Predator trait *v* linearly increases but not more than under 48 hours transfer intervals (Fig. 5b). The attack rate *a* displays interesting patterns, as it in most cases first decreases slightly, but returns to initial values at later stages (Fig. 5c). Again, conversion rate *b* linearly decreases but with little differences between coexistence times (Fig. 5c).

**Fig. 5.**
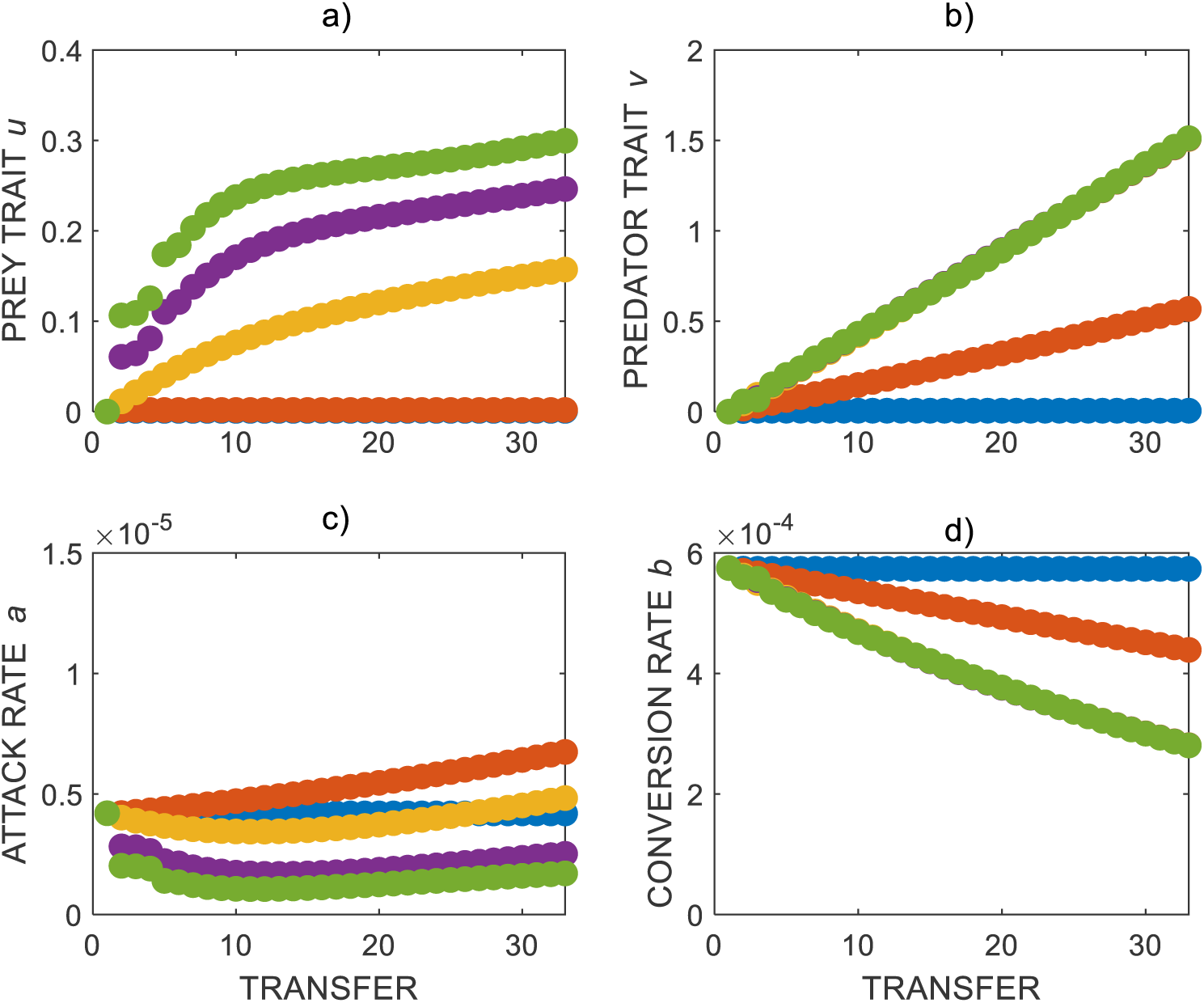
Evolutionary trajectories for bacteria and ciliates under different times of coexistence. a) prey trait *u*, b) predator trait *v*, c) predator attack rate *a* and d) predator conversion rate *b*. Dots in blue, red, yellow, magenta and green denote increasing times bacteria and ciliates grow together before next transfer (transfer interval 24h (blue), 36h (red), 48h (yellow), 60h (magenta) and 72h (green) hours). The transfer volume was kept constant at 1%. There were 33 transfers in total.

We were also interested how evolutionary dynamics are predicted under smaller modification of coexistence times. Increasing the time of coexistence only slightly (only 2-8 hours) has enhanced impact on evolutionary trajectories (Fig. S4). Notably, increasing the coexistence time only initially results in an increase in anti-predator traits in the bacteria, while predation traits evolve little different compared to the standard transfer interval of 48 hours.

### Interaction between mortality and coexistence time

Because we saw both, mortality and coexistence, to affect ecological and evolutionary dynamics individually we next asked how these two parameters interact. For example, a low mortality rate and a high coexistence time both result in increased evolutionary rates and we were interested if the effects are additive and evolutionary rates further increase or are dominant and no further change is observed. To explore this question, we simultaneously modified mortality and coexistence time in our model and tracked the dynamics.

Our model predicts an interaction between mortality (the transfer volume) and the time of coexistence (transfer interval). Increasing the transfer volume obviously decreases the mortality of both species, thus allowing them to survive better (Fig. 6). Beyond extinction conditions, bacterial and ciliate densities are rather independent from mortality and coexistence time. A simultaneous decrease in mortality and increase in coexistence times has little overall effect on densities and ecological dynamics are rather stable. Bacterial densities are predicted to be highest at lowest mortalities and longest coexistence times (Fig. 6a). Contrary to this, we see highest ciliate densities at long coexistence times, but at intermediate mortality rates (Fig. 6b).

**Fig. 6.**
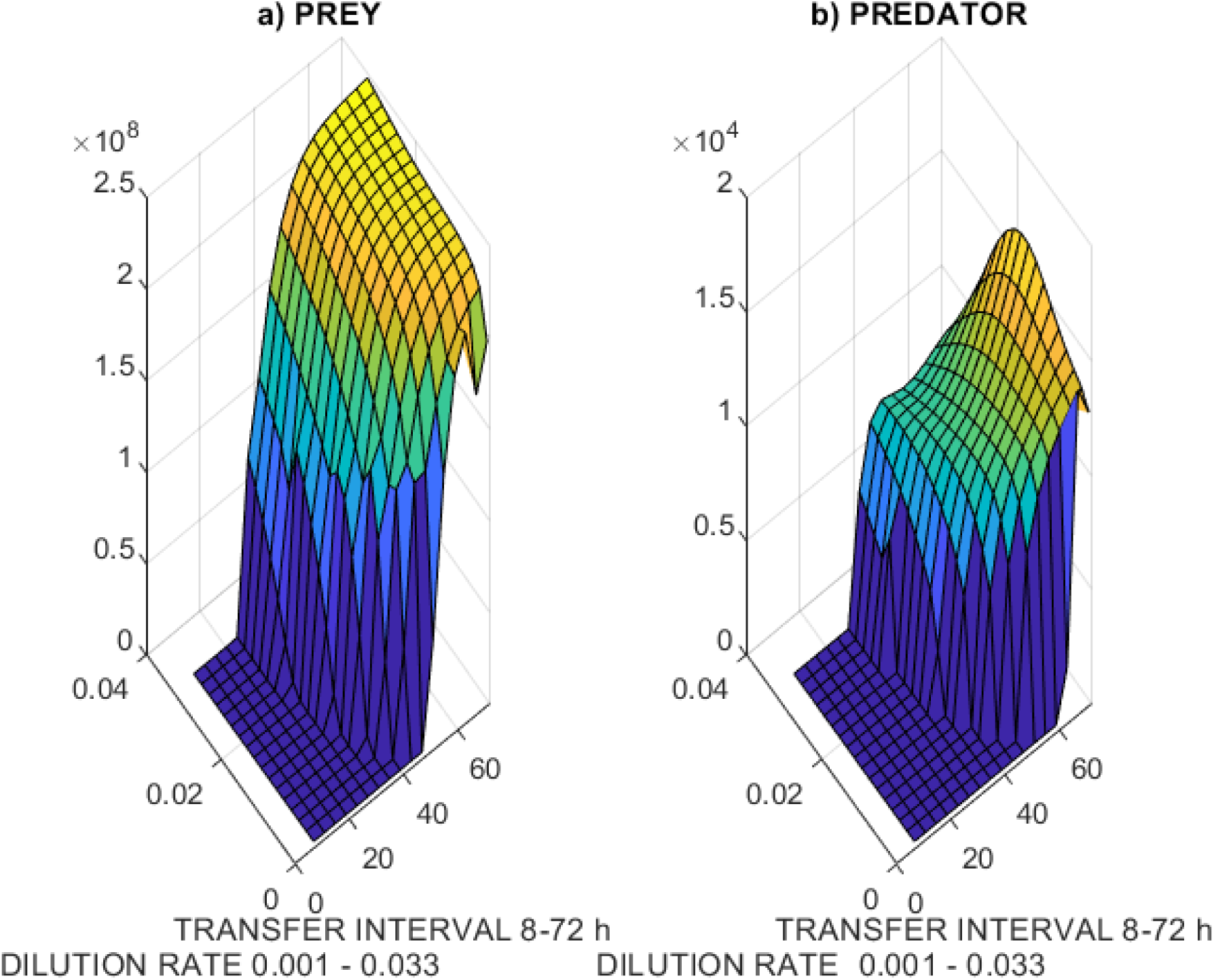
The combined effect of mortality and coexistence time on bacterial and ciliate densities. a) Bacterial densities across various transfer volumes and transfer intervals. b) Corresponding ciliate densities across the different parameters.

The evolutionary patterns, however, are predicted to be more depending in the interaction between mortality and coexistence time (Fig. 7). Bacterial anti-predator defence traits increase continuously and reach highest levels at maximum simulated coexistence times and lowest mortality rates (Fig. 7a). Please note, the evolutionary change seems to be more pronounced compared to the ecological change in density (Fig. 6a). For the ciliates we observe interesting evolutionary trajectories. While ecological maximum is predicted for intermediate mortality rates and long coexistence times (Fig. 6b), the predator trait *v* initially rapidly increases but suddenly plateaus off (Fig. 7b). Only at long coexistence times but very high mortalities there is a change in this trait again. The attack rate *a* display a curved mountain ridge pattern with a moving maximum so that the maximum attack rate is observed either under high mortality but long coexistence times, or under low mortalities but shortened coexistence times (Fig. 7c). The conversion rate *b* that was estimated for the naïve ciliates seems to be maladaptive and the model predicts constantly that conversion rates reduce independent of mortality and coexistence times (Fig. 7d).

**Fig. 7.**
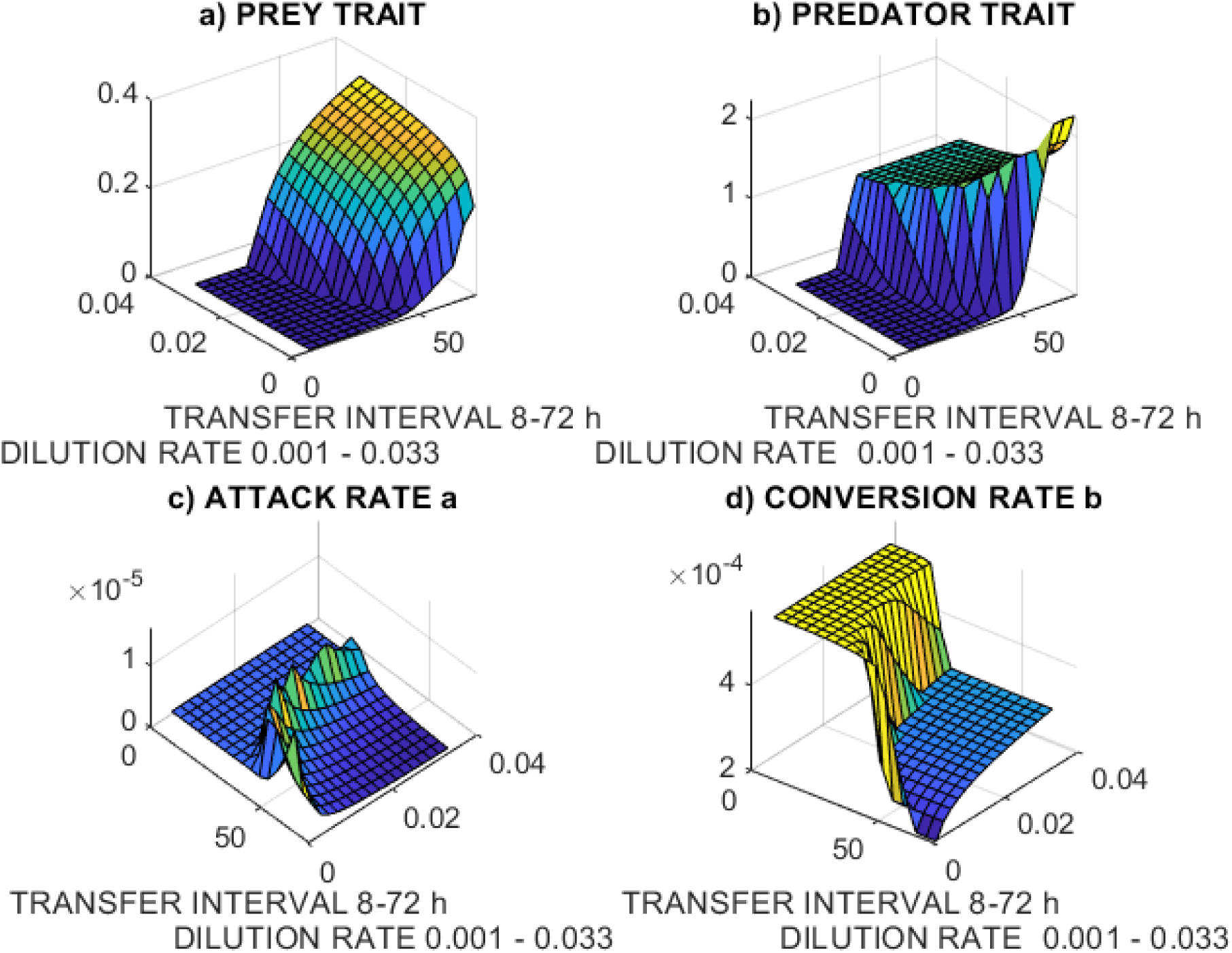
The interaction effect between mortality (transfer volume) and coexistence time (transfer interval) on evolutionary trajectories. Predicted evolutionary trajectories for anti-predator defence trait *u* of bacteria a), predator traits *v* b), the attack rate *a* c) and the conversion rate *b* d) of the ciliates.

### Sensitivity analysis and human caused impact

Our modelling approach offers additional insight how sensitive such a predator-prey experiment is related to protocol changes. In our model, transfer interval and transfer volume are always exact. However, after all, humans are not robots and mistakes can happen. Often there are slight changes in the protocol because researcher need to shift timing, e.g. occupied autoclave hasn’t finished in time, or pipettors work unprecise which remains unnoticed. To explore how a lack in precision affects the dynamics in such a system, we randomized parameters throughout the simulations. The first parameter we randomized was time of coexistence. For various reasons every researcher is aware, the transfer interval may deviate from the experimental protocol. So, what would be the effect if the protocol assumes starting a new growth cycle exactly after 48 hours with a transfer activity at noon 12 pm but the transfer happens any time between 9 am and 3 pm (Fig. S5a)? In this scenario, the ecological dynamics begin to display considerable variation (Fig. S5b). Particularly the predator densities fluctuate a lot. Surprisingly, the evolutionary trajectories seem to be rather robust for this type of variation (Fig. S5c).

Another parameter hard to control when starting the experiment is the effect of initial population densities added to the experiment. Researchers commonly estimate the densities of these microorganisms but of course the wanted densities can only roughly be added because of the miniature nature of the study system. To simulate this, we started our model assuming different initial densities for bacteria and ciliates. Differences in initial prey densities have little effect on ecological and evolutionary dynamics (Fig. S6). Increasing or decreasing the bacterial densities to start the experiment, is predicted to have no impact. Increasing the initial ciliate density also has little ecological and evolutionary effects (Fig. S7). Only the initial predator densities seem to be affected, but after a few growths cycles this initial effect should be lost.

Finally, right before transfer, populations may have patchy distribution which would result in variation of mortality rates when the community is not well mixed. We simulated variable mortality rates by randomizing the transfer volume (Fig. S8a). The result of this is again that ecological dynamics destabilize (Fig. S8b), however evolutionary dynamics are rather robust (Fig. S8c).

## Discussion

The question how communities change in a deteriorating environment is essential to predict future ecosystem functioning and services [27,28]. With progressing global change, acidification and nutrient enrichment of ecosystems and many other stressful factors, organism’s mortalities increase, and interaction networks may be disrupted. We used a mathematical model to simulate ecological and evolutionary dynamics of a life predator-prey system. We feel, the weakness of our approach making deductions only from model predications without further experimental validation turns into a strength as it allows us to explore many core parameters in fine detail. We would be unable to test all these different parameter settings in experiments but hope that this study spikes future experiments taking ideas up.

Our model predicts that ecological dynamics of experimental bacteria-ciliate communities are rather invariant for changes in mortality rate and coexistence time (Fig. 2, Fig. 4). The densities of bacteria and ciliates, which is the life predator-prey system we studied, only weakly depend on these parameters under regular transfer design. Only when mortality rates become too high or the coexistence time too short, which results in extinction, there are changes in population densities. Interestingly however, our model suggests that evolutionary dynamics are affected by these two parameters. Decreasing mortality rates are predicted to increase anti-predator defence evolution in the bacteria and attack efficiency in the ciliates, however in more complex ways for predators (Fig. 3). According with theory, the time of coexistence has also predicted effects, in the sense that longer coexistence times intensify evolutionary responses (Fig. 5).

Our model predictions are in line with previous findings demonstrating effects of increased mortality rates from abiotic change on community structures. Increased mortality rates caused by antibiotics affect ecological and evolutionary dynamics in this bacteria-ciliate system [14]. Similarly, competition, which also results in decreased population sizes of focal bacteria, interacts with predation and results in changed ecological and evolutionary dynamics [16]. Our finding that evolutionary trajectories are more affected than ecological dynamics is a bit in contrast with other studies, however. Increased mortality rates have been shown to result in community composition changes in bacterial communities [12], thus more on the ecological side. This study beautifully demonstrates that changes in mortality rates shift dominance of competing bacterial species from fast growers that are weak competitors towards slow growing species that are highly competitive. We expect this contradiction resulting from the different nature of the study system to explore bacterial competition rather than predator-prey interactions. Maybe it is also worth to mention that this study did not explore evolution, thus no inferences are possible. Our data are also in contrast with a different predator-prey system, namely rotifers grazing on algae. In this system increasing or decreasing mortality rates greatly impacts the nature of ecological interaction [8]. Changing the global mortality rate shifts the rotifer-algal densities between equilibrium and stable limit cycle states. However, this system follows a bit different experimental approach as there is a constant mortality rate because the organisms were grown in chemostat systems. In accordance with our study, the algal population quickly evolves in form of alternating genotype frequencies of contrasting defence level [29]. Other bacterial studies inducing high mortality rates at regular intervals also detect evolutionary changes in interaction [3,5,22], thus we think our findings represent a general pattern that evolution, and also ecology, depends on global mortality rate and the time species interact with each other.

The exact mechanisms why ecological dynamics are rather stable cannot fully be explained in our study. It is however likely that bacteria and ciliates reach the environmental imposed growth maximum quickly enough to result in stable dynamics. The original study [14] used nutrient limited medium which only supports low overall densities. Also, it is important to note that this system only reveals end points of each growth cycle, unlike a chemostat system, and inter-interval dynamics between growth cycles may be rather different resulting in similar final densities. The exact evolutionary mechanisms also remain open. Whereas evolutionary trajectories look rather clear for the bacteria and are well in line with theoretical predictions, the ciliate coevolution is less intuitive. First we have to emphasize that ciliate coevolution is also a model prediction [25], as looking for this was not scope of the original study [14]. That ciliates coevolve in such a system is however suggested by another study comparing naïve and potentially coevolved ciliates from the same system [16]. Still, why some predator traits evolve is difficult to explain and future studies will be needed to explore this in more detail.

Observing comparably little evolutionary change across settings in ciliates may depend in the fact that the underlying traits are depending on prey dynamics. This may be reflected by the equal ratios seen under different scenarios (Fig. S3). Perhaps evolutionary forces are reduced if ratios between bacteria and ciliates are little changing. The biological result makes sense as rate of evolution is expected to decline over time because of imposed costs, which need to be ameliorated before further change can happen.

Our findings support experimental approaches exploring ecological and evolutionary dynamics in microbial communities, suggesting these are good way to gain further insight into related questions, but call for caution in deciding for the experimental design. These experiments will detect ecological and evolutionary dynamics, only the magnitude may depend on the experimental design. We hope future researchers will take these ideas into account when designing upcoming experiments.

## Methods

We used a quantitative genetics approach and analysed the observed predator-prey interactions of this study applying a pure ecological Lotka-Volterra model [30] modified to explain co-evolution between the prey and predators [31,32]. Recall that the Lotka-Volterra model given as

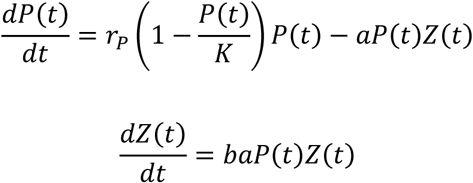

where *P* and *Z* denote the prey and predator populations, *r*_*P*_ is the prey growth rate, *K* is the carrying capacity, *a* is the attack rate and *b* is prey to predator conversion rate.

In the co-evolutionary version (Kaitala et al. 2019) the Lotka-Volterra model is revised such that the attack rate and the conversion rate are functions of auxiliary trait variables *u* and *v* of the prey and predator, respectively. The trait variables have dynamics of their own, the purpose of which is to maximize the fitness of the corresponding species. Thus, the coevolutionary model can be presented as follows

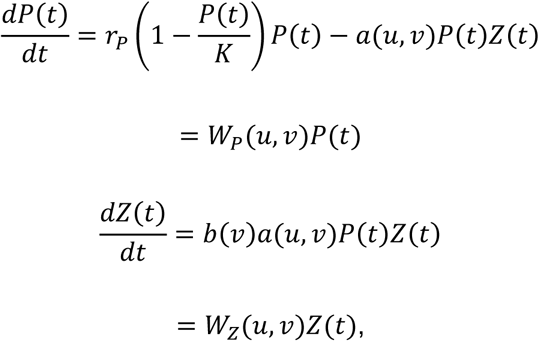

The trait dynamics are given as follows [31,33]

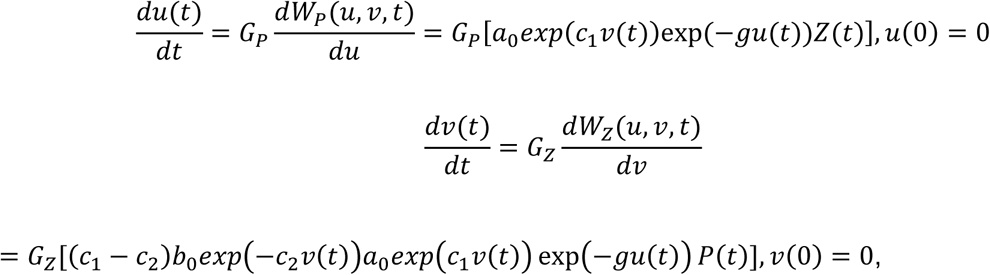

where *G*_*P*_ and *G*_*Z*_ are parameters determining the speed of the evolution of the traits. In the experimental data studied, the ancestral individuals in each species did not have any earlier history of occurring together in a predator-prey interaction. Thus, initial values of the traits *u* and *v* are chosen to be equal to 0. For more details about the model please see our previous study [25].

We next study effects of the changes in the experimental setting maintaining the original estimated model parameters [25] while modifying mainly the dilution rate (mortality) or growth period (transfer interval).

## Author contribution

VK and TS designed the study, VK wrote the mathematical model with input from TS and analysed the results, TS wrote the first draft of the manuscript.

## Data Availability Statement

All relevant data are within the paper and its Supporting Information files.

