## Supporting Material for "Mortality and coexistence time both cause changes in predator-prey co-evolutionary dynamics"

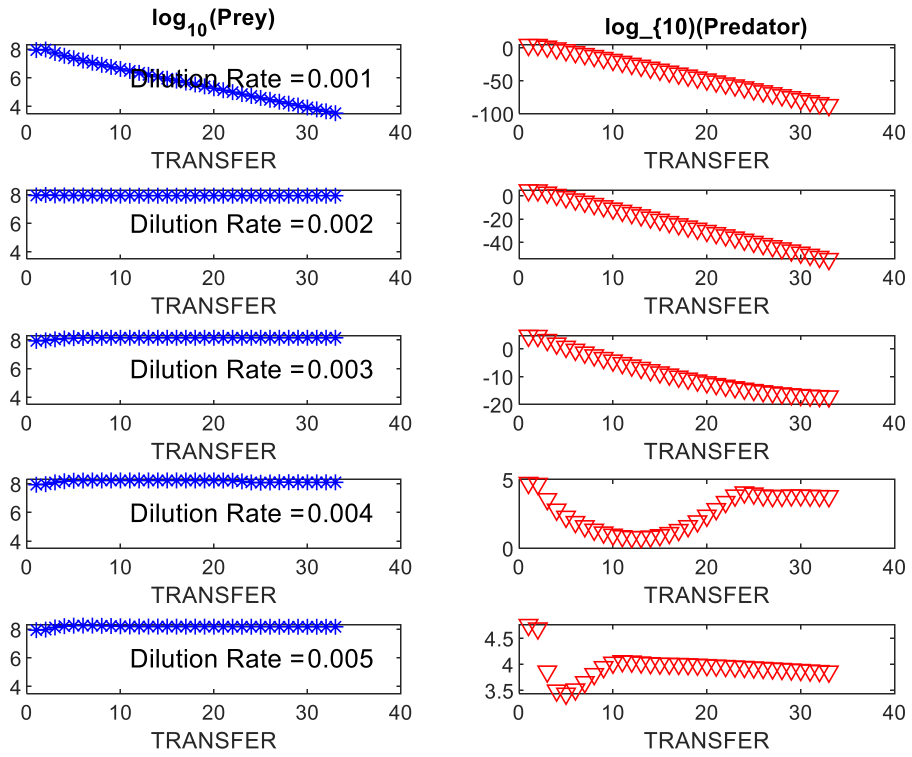


Figure S1. The effect of extremely high mortality rates at a transfer interval of 48 hour. The experiment is maintained for 33 transfers.

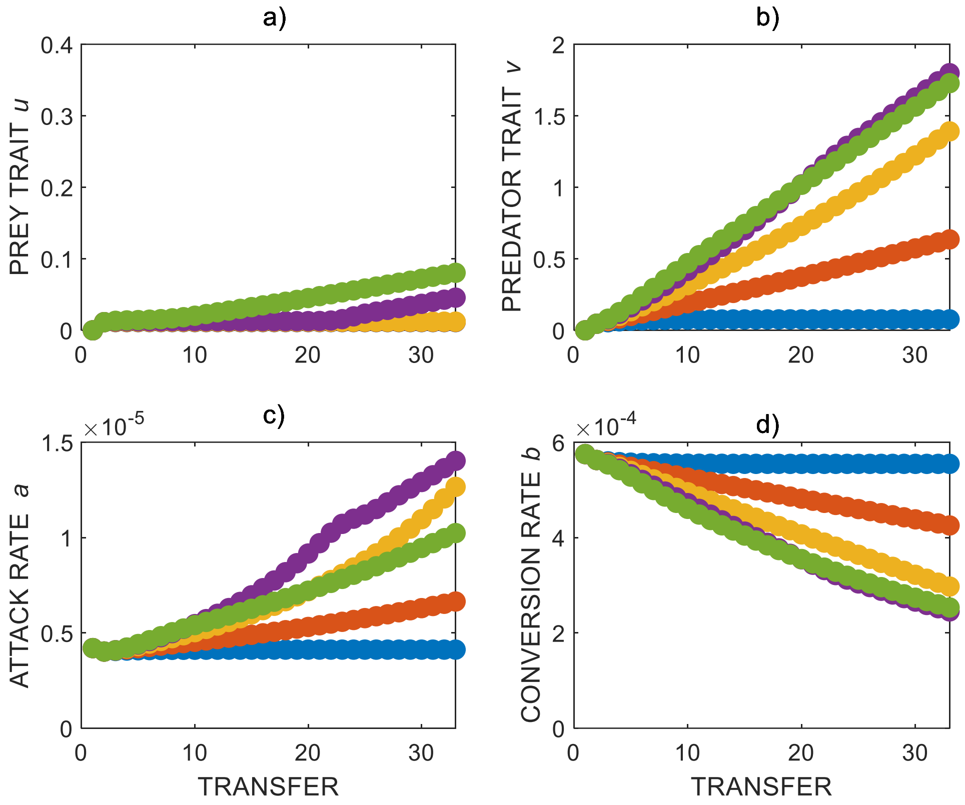


Fig. S2. Evolutionary trajectories under very high mortality rates. a) prey trait *u*, b) predator trait *v*, c) predator attack rate *a* and d) predator conversion rate *b*. Squares in blue, red, yellow, magenta and green denote increasing mortality rates respectively (dilution rate).

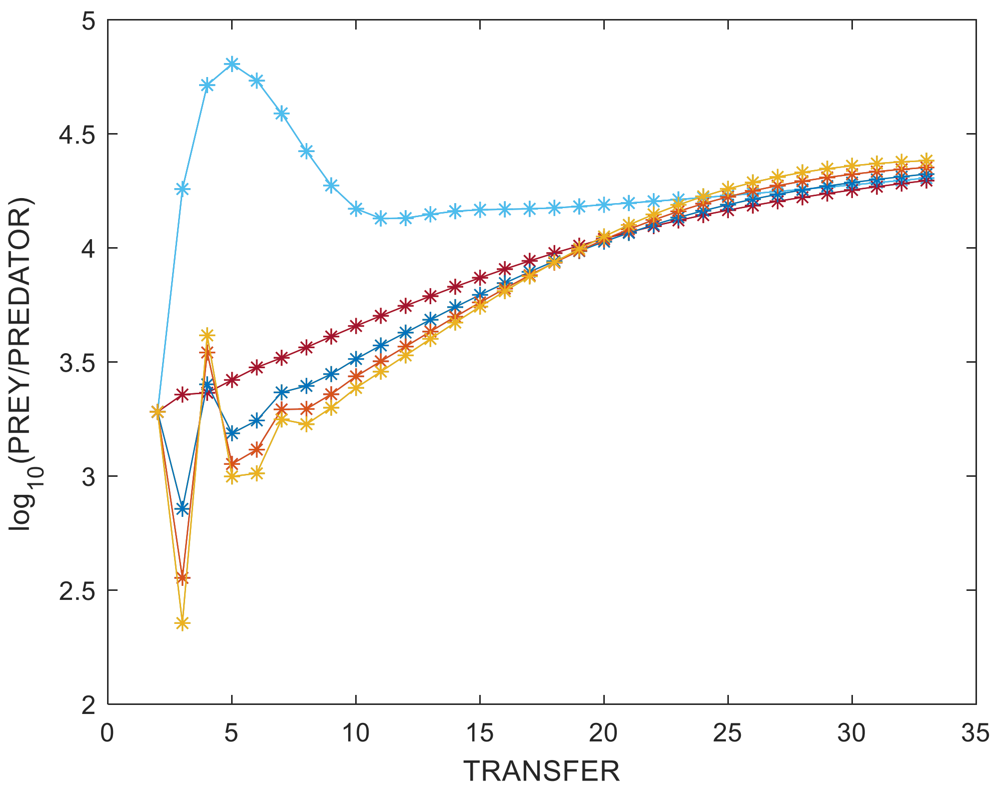


Fig. S3. Level of coexistence between bacteria and ciliates. Blue denotes very high mortality and green the lowest mortality level.

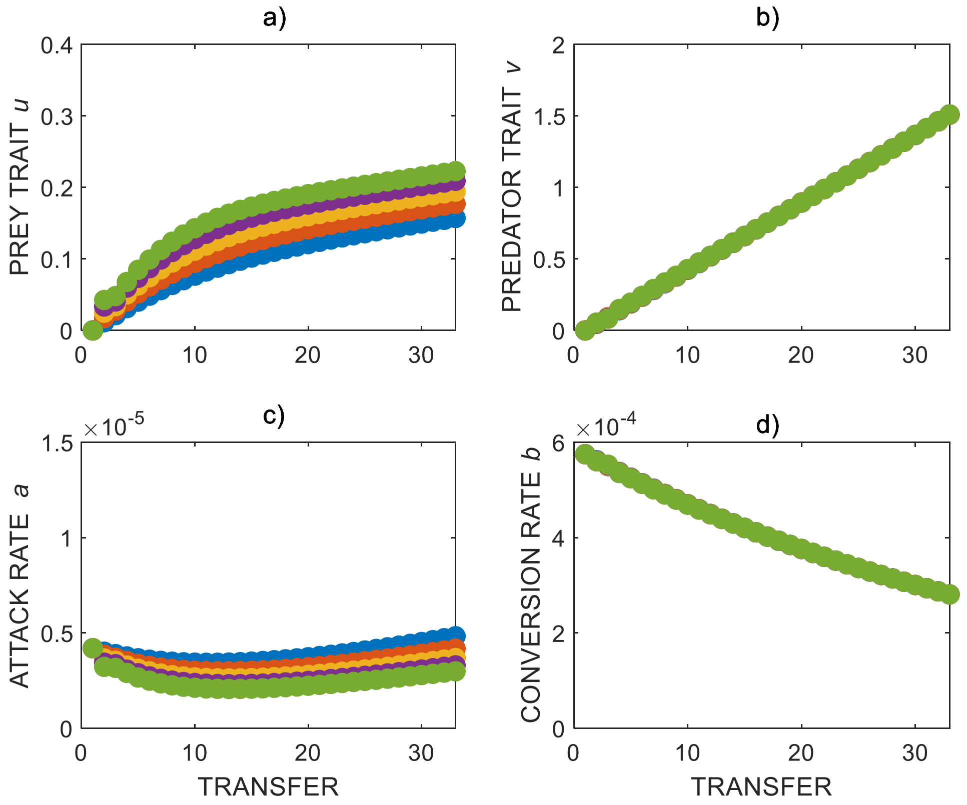


Fig. S4. Evolutionary trajectories for slightly increased times of coexistence. a) prey trait *u*, b) predator trait *v*, c) predator attack rate *a* and d) predator conversion rate *b*. Dots in blue, red, yellow, magenta and green denote increasing coexistence times (transfer intervals). Here the transfer intervals were increased by 2 hours from 48 to 56 hours.

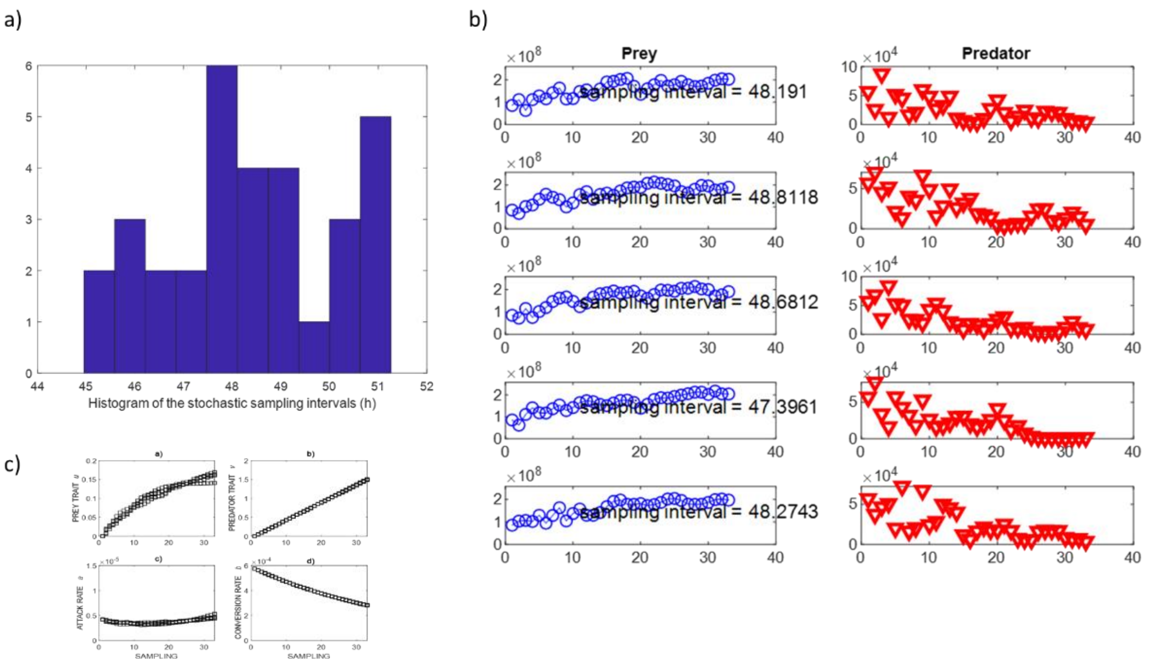


Fig. S5. The effect of variation in transfer intervals. a) Actual transfer happens between 45 and 51 hours and not exactly after 48 hours. b) The ecological dynamics for bacteria (blue) and ciliates (red). c) evolutionary trajectories. Five **replicates** of the experiments with stochastic sampling intervals (T2=48 + (rand-0.5)×3). The expected value is 48 hours. The sampling intervals are independent. The model used a dilution rate of 1%.

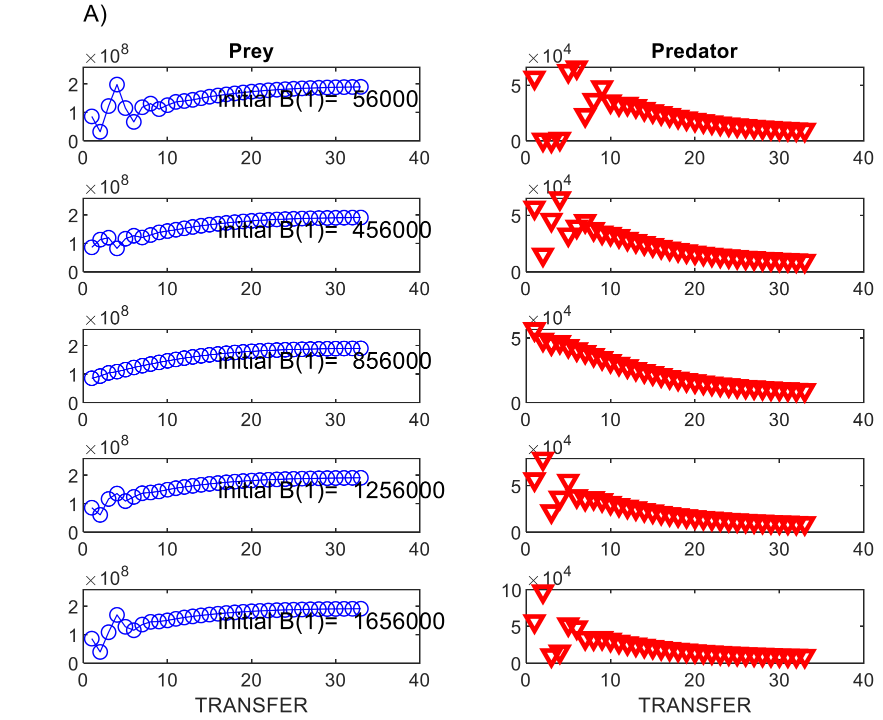


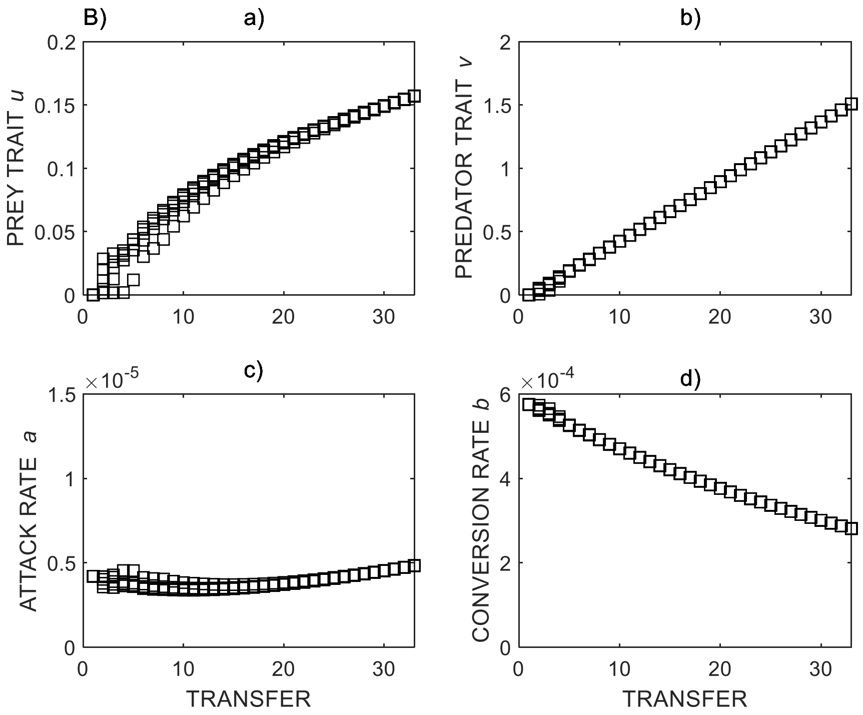


Fig. S6. The effect of different initial bacteria concentrations. A) Ecological dynamics. B) Evolutionary dynamics. The model used a transfer interval of 48 hours and a dilution rate of 1%.

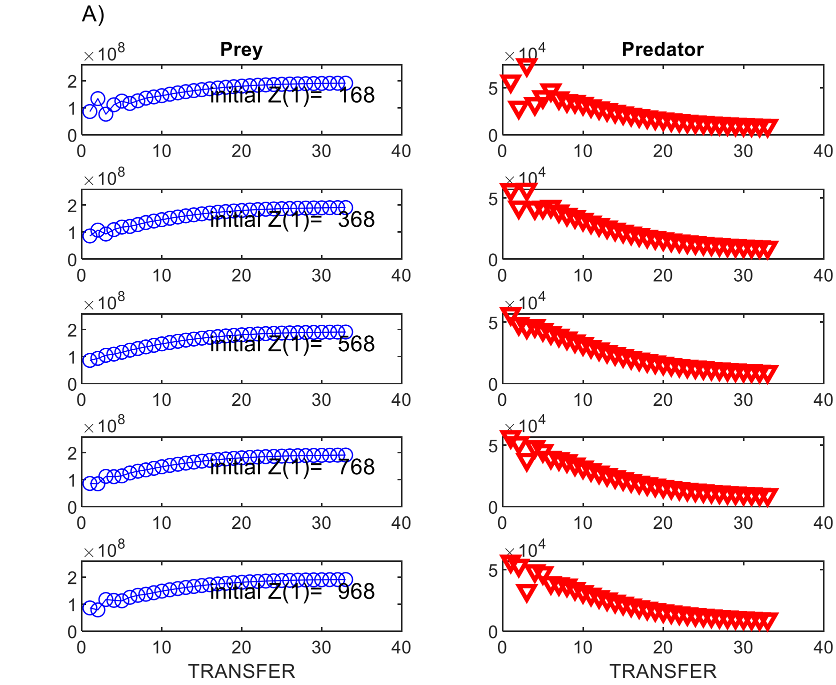


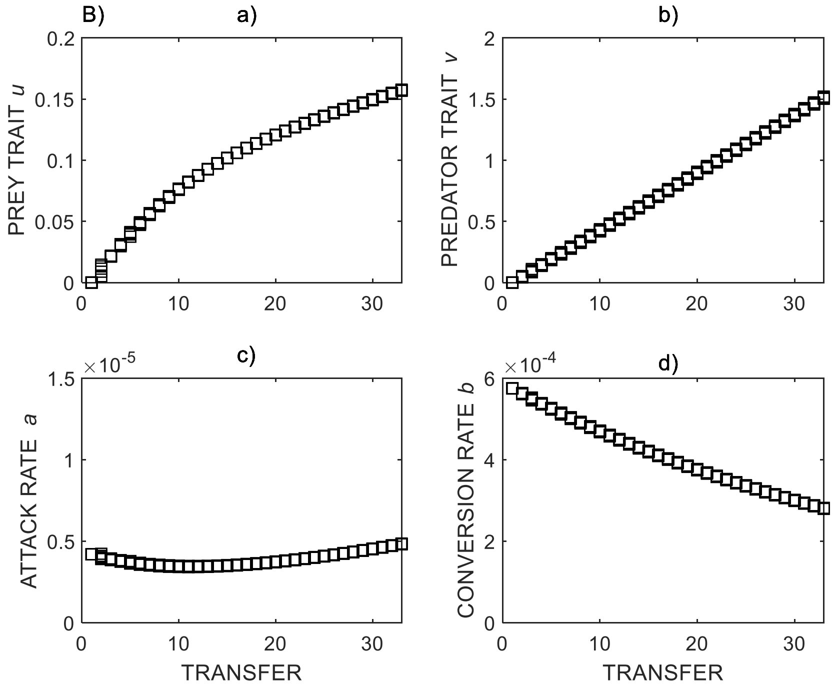


Fig. S7. The effect of different initial ciliate concentrations. A) Ecological dynamics. B) Evolutionary dynamics. The model used a transfer interval of 48 hours and a dilution rate of 1%.

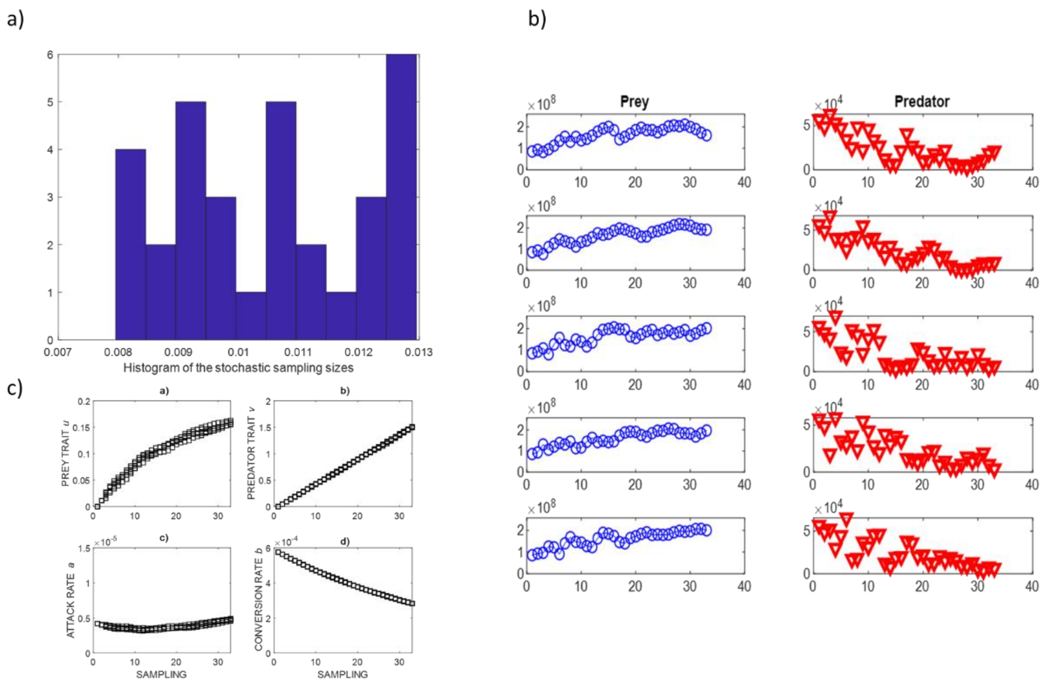


Fig. S8. The effect of different mortality rates implemented by randomized sampling volumes. a) Randomization of dilution rates. b) Bacterial and ciliate densities. c) Evolutionary rates. The model used a transfer interval of 48 hours and a dilution rate of 1%.
